# Trivial Excitation Energy Transfer to Carotenoids is an Unlikely Mechanism for Non-Photochemical Quenching in LHCII

**DOI:** 10.1101/2021.10.18.464810

**Authors:** Callum Gray, Tiejun Wei, Tomáš Polívka, Vangelis Daskalakis, Christopher D. P. Duffy

## Abstract

Higher plants defend themselves from bursts of intense light via the mechanism of Non-Photochemical Quenching (NPQ). It involves the Photosystem II (PSII) antenna protein (LHCII) adopting a conformation that favours excitation quenching. In recent years several structural models have suggested that quenching proceeds via energy transfer to the optically forbidden and short-lived S_1_ states of a carotenoid. It was proposed that this pathway was controlled by subtle changes in the relative orientation of a small number of pigments. However, quantum chemical calculations of S_1_ properties are not trivial and therefore its energy, oscillator strength and lifetime are treated as rather loose parameters. Moreover, the models were based either on a single LHCII crystal structure or Molecular Dynamics (MD trajectories) about a single minimum. Here we try and address these limitations by parameterizing the vibronic structure and relaxation dynamics of lutein in terms of observable quantities, namely linear absorption (LA) transient absorption (TA) and two-photon excitation (TPE) spectra. We also analyze a number of minima taken from an exhaustive meta-dynamical search of the LHCII potential energy surface. We show that trivial, Coulomb-mediated energy transfer to S_1_ is an unlikely quenching mechanism. Pigment movements are insufficient to switch the system between quenched and unquenched states. Modulation of S_1_ energy level as a quenching switch is similarly unlikely. Moreover, the quenching predicted by previous models is likely an artefact of quantum chemical over-estimation of S_1_ oscillator strength and the real mechanism likely involves non-trivial inter-molecular states.

## 1 Introduction

Non-photochemical quenching (NPQ) in higher plants is a regulatory response to a sudden increase in light intensity [1, 2, 3, 4]. It is a (mostly [5]) reversible down-regulation of the quantum efficiency of the Photosystem II (PSII) light-harvesting antenna (LHCII) with the purpose of defending the saturated reaction centres from over-excitation and photoinhibition [6, 7]. Essentially, it is due to the creation of exciton-quenching species within LHCII which trap and dissipate chlorophyll excitation before it can accumulate in PSII and damage the reaction centres. While the fine molecular details of the mechanism are still unclear, a general consensus has emerged over the basic scheme. It’s primary trigger is a acidification of the thylakoid lumen (ΔpH) due to a high rate of electron transport [8], in large part cyclic electron flow about PSI [9]. The ΔpH activate three NPQ components: the PSII antenna sub-unit PsbS [10], the enzyme violaxanthin de-epoxidase (VDE) [11], and the LHCII antenna proteins themselves [12, 13, 14]. VDE converts the violxanthin pool to zeaxanthin which may lead to violaxanthin-zeaxanthin exchange in the loose, peripheral xanthophyll-binding site of LHCII [15]. It has been shown that the presence of zeaxanthin affects the kinetics and amplitude of NPQ but is not a strict requirement for it [16, 17]. Similarly, it was shown that quenching can be achieved in the absence of PsbS if ΔpH is driven to non-physiological levels [18]. Either way (and the reader is directed to a comprehensive review of this complex and on-going topic [19]) the combined effect is to induce an in-membrane aggregation or clustering of LHCII [20] and some subtle internal conformational changes [21]. These somehow modulate the pigment-pigment and pigment-protein couplings to create a quenching species, although the nature of the quencher and molecular dynamics of the conformational ‘switch’ are still unclear.

In recent has become broadly (though by no means universally) accepted that the quencher is or involves one of the LHCII carotenoid (Cart) pigments [22]. These are attractive candidates as they are intrinsically quenched pigments, possessing a very short (≈10 ps) excitation lifetime relative to chlorophyll (Chl, ≈4-6 ns). Various mechanisms have been suggested such as: excitation energy transfer (EET) to the Cart which quenches simply by virtue of its short lifetime [23], mixing of the Chl and Cart lifetimes brought on by excitonic resonance [24, 25], and formation of fast-relaxing Chl-Cart CT states [26, 27]. With regard to which carotenoid, the lutein (Lut) bound to the L1 binding site [28] of the LHCII trimer is often cited [23] but zeaxanthin at an equivalent site in one of the minor PSII antennae has also been proposed [26]. We also note that Holzwarth and co-workers present a Cart-independent quencher model that involves Chl-Chl CT states [29, 30]. The differences between these different models often comes down to specific interpretations of highly-congested time-resolved spectral measurements on these complexes. Moreover, any involvement of the Carts is obscured by the fact that their lowest singlet excitation, S_1_, is optically forbidden and decays very quickly [31].

The X-ray structure of LHCII [32] can provide some insight into the quencher, particularly since it was found to correspond to a highly dissipative configuration, meaning it could serve as a model structure for the quenched state [33]. Several detailed models of this structure very accurately predicted the steady state and time-resolved spectra of LHCII [34, 35, 36] but they did not capture the dissipative character (in fairness that was never their goal). One possible reason for this was their neglect of the Carts, due to the fact that they contribute nothing to the spectrum in the red region of the spectrum and that there are no truly reliable methods for calculating the excitation energy and one-electron transition density of the S_1_ state. The latter is due to the strong electron correlations giving it a complex multi-electron character [37, 38]. Beginning in 2013 Duffy and co-workers used a semi-empirical quantum chemistry method to estimate the S_1_ transition density and its potential affect on the excitation lifetime of the LHCII crystal structure [39, 40, 41, 42]. These models suggested that quenching was due to EET from the Chl Q_y_ band to the S_1_ state of the centrally bound Luts (L1 and L2), followed by fast decay of S_1_. This EET was mediated by weak resonance couplings between Q_y_ and S_1_ (due to the latter’s lack of oscillator strength) and was therefore assumed to be incoherent (Förster transfer) and relatively slow (20-50 ps) relative to excitation equilibration across the Chls (≈1-2 ps). This is essentially the mechanism proposed by Ruban et al. based on global target analysis of transient absorption (TA) measurements on LHCII aggregate, although they propose L1 is the sole quencher [23]. Of course these models are all based on a single structural snap-shot and a highly artificial one at that. It therefore tells us nothing about how such quenching is switched on and off and can only very tentatively be applied to the actual *in vivo* quenching mechanism. More recently, the model was extended to a molecular dynamics simulation of the LHCII trimer within a lipid bilayer [43]. Although, a stable, unquenched conformation was not identified, it predicted that the Q_y_ - S_1_ coupling was highly-sensitive to very small changes in inter-pigment orientations, suggesting that the lifetime could be modulated by very subtle conformational changes. Unfortunately, this appears to have been incoherent for two reasons. Firstly, the coupling sensitivity appears to have been an artifact of the semi-empirical Hamiltonian used to calculate S_1_. Khokhlov and Belov showed that this sensitivity disappears when chemically accurate methods are used to calculate the S_1_ transition density [44]. Moreover, by simulating the near-identical CP29, Lapillo et al. showed that even with the semi-empirical method, large lifetime fluctuations are significantly dampened if one accounts for the excitonic structure of the Chl manifold in the complex (the previous model assumed a Chl-Lut dimer embedded in some coarse-grained, iso-energetic Chl pool) [45]. In addition to these errors, the model has a series of weaknesses that here we attempt to address:-

- ***The* S_1_ *excitation energy*** is neither easy to measure directly or calculate. Transient absorption in near-IR gives a phononless excitation energy of 14, 050 ± 300cm^−1^ for Lut in recombinant LHCII [46], while two-photo excitation (TPE) in native LHCII gave < 15, 300cm^−1^ [47]. the latter value is likely the first vibronic peak which is ≈ 1100cm^−1^ higher than the phononless peak, meaning the two values reasonably agree. Nevertheless, it is often treated like a free parameter and large changes its value have been proposed as a part of the NPQ switch [25, 45].
- ***The vibronic structure and relaxation dynamics of* S_1_** were not properly considered. It was treated as a single optical transition with a line-broadening function chosen to provide a convincing visual fit to the TPE of S_1_, which implied very large reorganization energies. It was assumed that reorganization on S_1_ was instantaneous and that inter-conversion (IC) to the ground state (S_0_) occurred with a single rate constant of 10 - 20ps [46]. The end result is a picture of S_1_ as an deep, irreversible trap. In reality, S_1_ is composed of several vibronic transitions that could couple differently to Q_y_, relax on finite timescales and undergo IC at different rates.
- ***Limited sampling of the LHCII potential energy surface (PES)*** means we might not be probing biologically relevant conformations. Unsteered molecular dynamics (MD) simulations start from a quenched minimum close to the crystal structure. Single molecule spectroscopy has shown that LHCII trimers will spontaneously switch between quenched and unquenched states but the typical dwell time in each is of the order 1-10 s [48], meaning prohibitively long unsteered simulations may be needed to capture this switching.

Here we attempt to correct these errors in several ways. We obtain a detailed picture of the S_1_ energy gap, vibronic structure and relaxation kinetics by fitting a detailed model to the TA kinetics of Lut in pyridine. These parameters (along with a secular Redfield model of the Chl manifold [49]) are then used to model energy relaxation in LHCII. The LHCII model structures that we use come from an exhaustive steered search of the LHCII PES which was previously published [50]. The motivation is to determine whether NPQ can realistically be switched on and off simply by altering the relative distance/orientation of Lut and its neighbouring chlorophylls.

## 2 Results

### 2.1 Steady-state spectra of the chlorophyll excitonic manifold

The Chl-Chl relaxation dynamics are modelled according to the method in [49] and briefly recapped in the Methods. For a given LHCII monomer trajectory we take a set of uncorrelated snap-shots and for each calculate the population relaxation. The snap-shots sample disorder in the inter-pigment excitonic couplings and the different minima in the original steered MD may reveal differences in the *average* couplings. We do not calculate the Chl excitation (site) energies but simply take the average values reported in [35]. The reason for this is partly to spare computational expense and partly because these fluctuations have almost no effect on quenching [52]. To check the validity of the model we calculate the linear absorption (LA) and fluorescence (FL) profiles, examples of which are shown in Figs. 1a and 1b, adding Gaussian disorder to the site energies to reproduce the homogeneous broadening.

**Figure 1:**
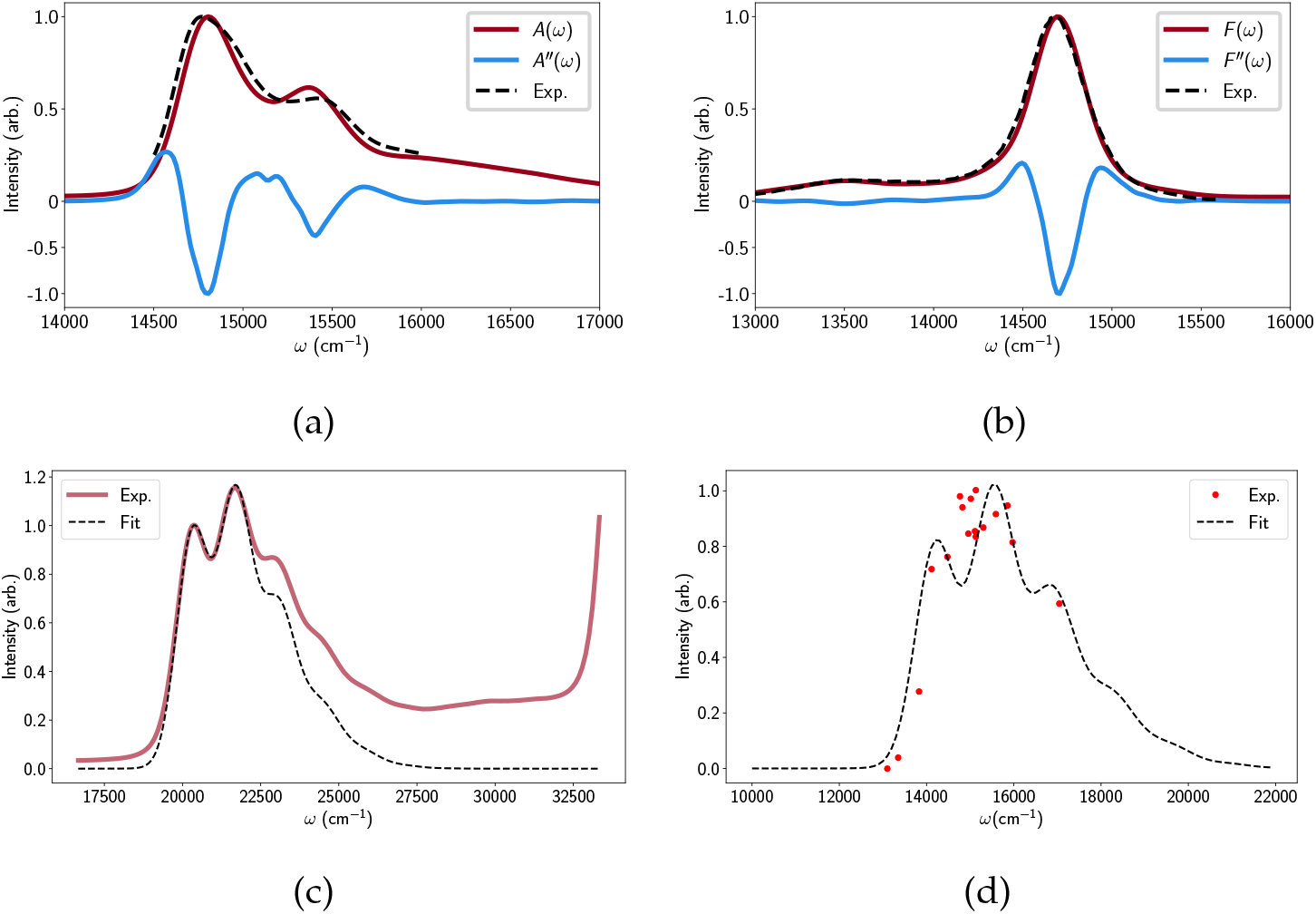
(a) A calculated linear absorption spectrum derived from one of the minima (red line) compared to the experimental spectrum (dashed line) [48]. The second derivative of the calculated absorption is shown (blue line) to highlight the Chl a and b peaks. The calculated spectra were essentially identical for all LHCII minima probed. (b) Calculated and experimental fluorescence profiles. (c) The calculated (dashed line) and measured linear absorption spectra of Lut in pyridine. (d). The calculated (dashed line) and measured (red dots) [51] TPE spectrum of Lut. All calculation parameters were taken from the TA fit apart from a 150cm^−1^ blue-shift to account for the fact that the TPE measurements were performed in octanol

### 2.2 Relaxation kinetics of lutein in pyridine

For Lut we adopt the Vibrational Energy Relaxation Approach (VERA) [53, 54] to reproduce several independent spectral measurements. The details are discussed in Methods (and Section C of the Supporting Information) but essentially the four singlet electronic states (|S_0_〉, |S_1_〉, |S_2_〉, |S_n_〉) are replaced by sets of vibronic states, (|i_a_1_a_2__〉 = |i〉 |a_1_〉^i^ |a_2_〉^i^), where i is the electronic index and a_1_ and a_2_ are the vibrational quantum numbers associated with the high-frequency, optically-coupled C - C and C = C modes respectively [53]. The LA is given by the sum of all vibronic transitions belonging to |S_0_〉 → |S_2_〉 (weighted by the Franck-Condon overlaps and the initial populations on |S_0_〉) and is shown in Fig. 1c alongside the experimental profile for Lut in pyridine. The fit is very good up to the blue edge of the first vibronic peak after which there is a deviation due to contributions from different geometrical conformers that are not accounted for in our model [55]. The rise after 27, 500cm^−1^ is a solvent artefact.

The static properties of S_1_ and all dynamical properties were obtained by fitting the VERA model to the TA of Lut in pyridine. Figs. 2a and 2b show the calculated and experimental difference spectra at intermediate (1 – 20ps) and long (10 – 42ps) delay times respectively. The sub-ps kinetics are not shown as they are less relevant to the final quenching model. The S_1_ Excited State Absorption (ESA, positive feature around 18, 000cm^−1^) is well fit but there is some discrepancy for the Ground State Bleach (GSB, negative feature) at earlier times. While the fit can be improved by adjusting the S_2_ parameters, this disrupts the original LA fit. This could be linked to the GSB-distorting local heating effects that were previously reported [54] or simply an artefact. Either way it is the S_1_ parameters and kinetics that we are primarily interested in. All fitting parameters are reported in Section A of the Supporting Information but there are a few key quantities:-

- **The phononless S_1_ energy**, ε_S_1__ = 14, 050cm^−1^: We assumed the previously reported value during the fit to reduce the number of free parameters. Varying ε_S_1__ naturally ruins the fit but it can be recovered by adjusting other parameters (mainly ε_S_n__ and the dimensionless displacements between S_1_ and S_0_. As an independent check we calculated the S_0_ → S_1_ line-shape and compared it to the TPE of Lut in octanol [51]. This is shown in Fig. 1d and apart from a 150cm^−1^ blue shift to account for the different solvent there is a reasonable visual agreement. However, we must note that the fit (nor the data, really) does not match the mirror image of the S_1_ FL line-shapes observed for Carts such as neurosporene and spheroidene [56]. These have a much less defined vibronic structure and deconvolution suggests that the largest peak is the 0 - 2 line (|0_00_〉 → |1_01_〉 in our model) rather than 0 – 1. We found it impossible to reproduce such a line-shape while retaining any kind of fit to the TA and TPE data. This may be a limit of the displaced oscillator model but it was later suggested that S_1_ FL measurements may be distorted by the presence of cis-isomers [57].
- **The S_1_ lifetime**, 〈τ_S_1_→S_0__〉 ≈ 14ps: Fig. 2c shows the evolution of total population on S_2_, S_1_ and S_0_. S_2_ → S_1_ inter-conversion (IC) occurs on the 100fs timescale while S_1_ undergoes near mono-exponential decay in within the the 10 – 20ps range usually quoted for xanthophylls.
- **Vibrational relaxation on S_1_**, 〈τ_vib-S_1__〉 ≈ 1ps: Fig. 2d shows the population evolution of the S_1_ vibronic levels |1_00_〉, |1_10_〉 and |1_01_〉. While it is difficult to assign a single lifetime to a multi-component, transient process it is clear that vibrational relaxation on S_1_ is an order of magnitude faster than S_1_ IC but not instantaneous.
- **Vibrational relaxation on S_0_**, 〈τ_vib-S_0__〉 < 14ps: There is a very small transient population on |0_01_〉 and |0_10_〉 which reaches a peak at 9ps (not shown) and makes a very small contribution to the blue shoulder on the S_1_ ESA. However, unlike Carts such as canthaxanthin and rhodoxanthin [54], the S_1_ IC is too slow to generate a vibrational population inversion on S_0_ and hence there is no S*-type signal. [54]

**Figure 2:**
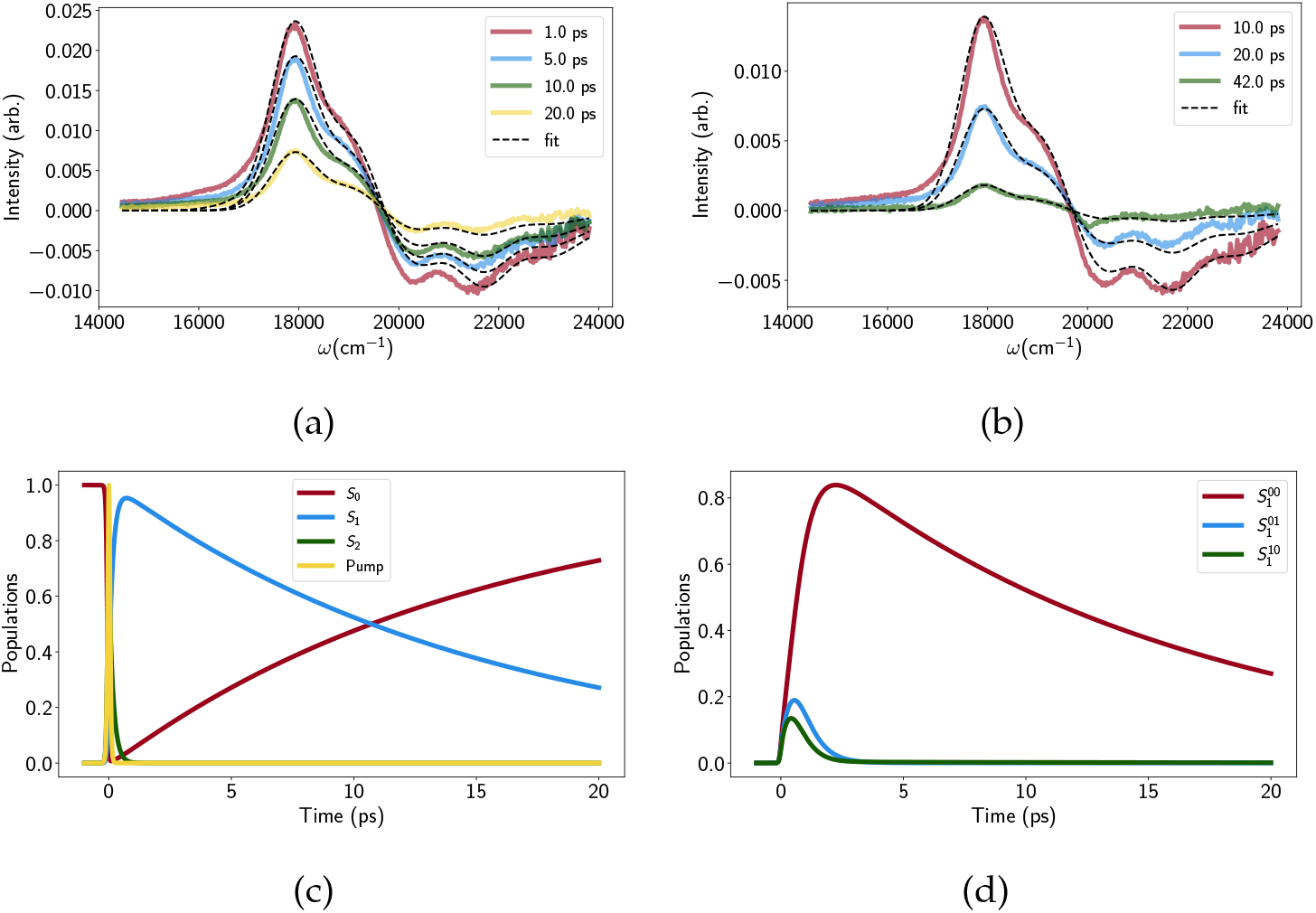
Transient absorption traces for (a) medium times and (b) long times, with experimental data shown in solid, coloured lines and model fits shown as dashed lines. (c) The evolution of the total population on each electronic state as a function of time, along with the temporal shape of the pump pulse. Note that a large part of the S_2_ → S_1_ decay (green line) overlaps with the pump. (d) Population evolution of the 3 lowest vibrational levels on S_1_.

### 2.3 Excitonic states and intermolecular couplings

As a baseline we first simulated relaxation in the LHCII crystal structure [32] following protonation and minimization. The Chl-Chl couplings are essentially as previously reported[34, 35] and diagonalization leads to a set of exciton states that have already been describe elsewhere [34]. Briefly, in the range 15, 200 - 15, 500cm^−1^ are a set of almost single-molecule (unmixed) Chl *b* states. Within 14, 700 - 15, 200cm^−1^ are a excitonic state typically localized on dimers or trimers of Chl *a*. Particularly relevant to NPQ is the terminal emitter state at around 14, 730cm^−1^ which is localized on the Chl *a*610-*a*611-*a*612 domain closely associated with Lut1, which we label |TE^−^〉. There is also the high-energy ‘anti-bonding’ equivalent, |TE^+^〉, at around 15, 120cm^−1^. This is shown diagrammatically in the Fig. 3.

**Figure 3:**
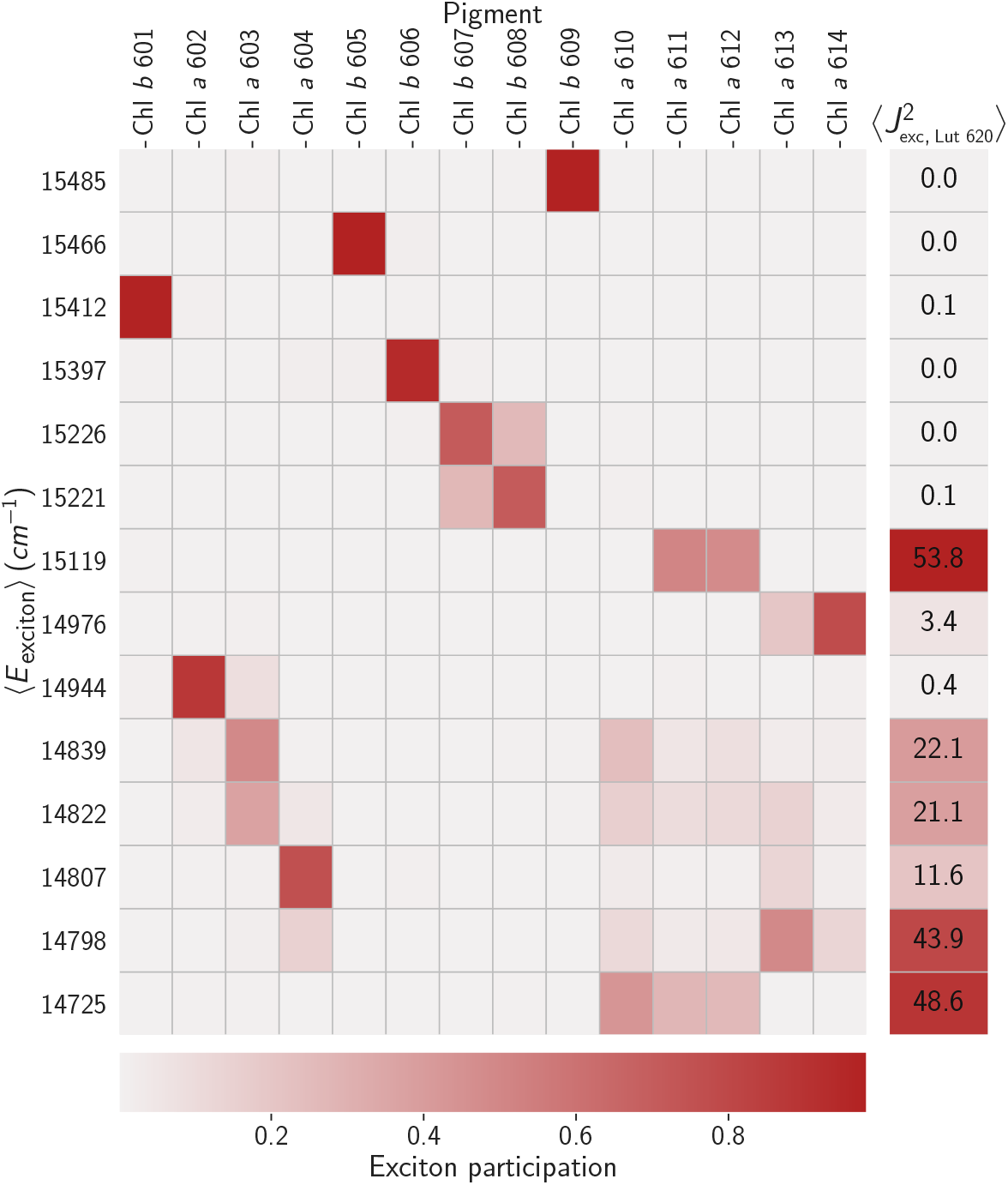
The grid shows the average energies, 〈E_i_〉, of the excitonic states and the average exciton participations, 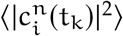, for a typical minimum. The right-most column lists typical average values of the square couplings between the exciton states and the 0 - 0 transition on Lut. The coupling to higher vibronic transitions are simply weighted by the relevant Franck-Condon overlaps.

We then consider the purely electronic Chl-Lut1 couplings, 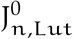, in the site basis we have the same picture as previously reported [40], weak (10 - 20cm^−1^) couplings to the terminal emitter Chls and negligible couplings otherwise. Excitonic mixing among the Chls naturally mixes these couplings, the strongest 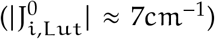 being between Lut1 and |TE^−^〉 and |TE^+^〉. These small couplings fully justify [52] our mixed kinetic model in which the Chl excitonic and Lut1 vibronic subsystems can exchange energy incoherently.

When we model the relaxation kinetics the presence of Lut1 results in a decreased excitation lifetime of τ_ex_ ≈ 500ps, compared to the unquenched value of 4ns [33]. The pathway is two-fold, involving fairly-reversible transfer from |TE^+^〉 to the near-resonant 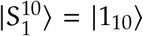 level and steep down-hill transfer from |TE^−^〉 to 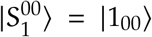 (see Fig. 4a). The transfer is typically slow. For example the rate constant for transfer from |TE^−^〉 to 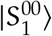 is 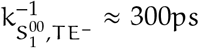. There are, however, several pathways that contribute, involving other exciton states and the 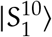 vibronic level, resulting is a net timescale of ≈ 100ps. This is too slow for any transient accumulation of population on 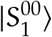 (see Fig. 4b).

**Figure 4:**
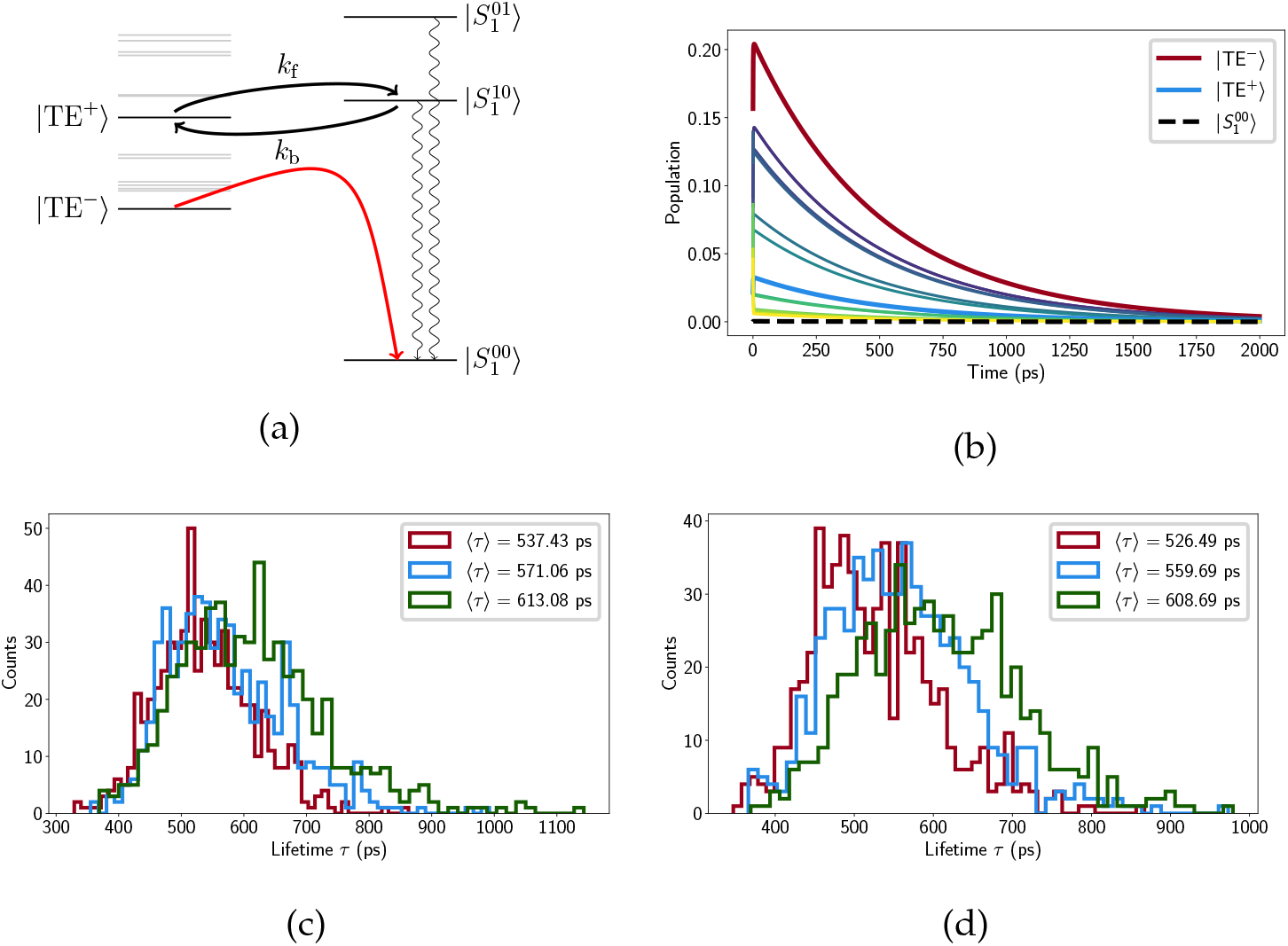
(a) Schematic representation of our model with the Chl excitonic manifold shown on the left and relevant carotenoid vibronic states on the right (energies to scale). (b) Chl exciton populations calculated from the crystal structure as a function of time. The exciton states associated with the terminal emitter labelled, along with the population on lowest vibrational level of 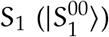. Selected histograms of the mean excitation time 〈τ〉 of different LHCII minima (from the steered MD) for low ((c)) and neutral pH (d) respectively.

While these results are essentially identical to those previously reported [40], it is important to realize that the absolute value of τ_ex_ is not meaningful, due to the fact that the Chl-Lut couplings are derived from un-scaled S_1_ transition charges from quantum chemical calculations [44].

### 2.4 Exploring the LHCII potential energy surface

We calculated the average, *relative* mean excitation times for several minima identified by a previous steered MD study [50]. It was reported that different monomers within the same LHCII trimer could access different conformational states and so we consider the monomer in our calculations. The minima are broadly classified into ‘low pH’ and ‘neutral pH’ depending on the protonation state of state of several lumen-exposed residues. In all minima there were fluctuations in the *snapshot lifetimes*, 300 < τ_ex_ < 1000ps, but the average value varies little within the range 500 < 〈τ_ex_〉 < 600ps (see Figs. 4c and 4d).

It is premature to say that ‘all of these minima are quenched’ but we can state that there is no evidence of a simple, purely-geometric switch between states with significantly different lifetimes. We find (as previously noted [41]) that τ_ex_ is correlated with Lut1-Chl *a*612 coupling but the coupling is not sufficiently sensitive to the small movements of the pigments to produce any kind of functional transition.

### 2.5 S_1_ excitation energy and asymmetry between Lut1 and Lut2

If we trust the TA-derived value of ε_S_1__ = 14, 050cm^−i^ then the relative arrangement of excitonic and vibronic levels is notable (see Fig. 4a). |TE^+^〉 and 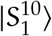 are near-resonance but since |TE^+^〉 acquires little exciton population at room temperature (see Fig. 4b) this is not a very effective pathway for quenching. The terminal emitter state, |TE^−^〉, lies almost precisely in the middle of 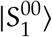 and 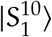 meaning any reasonable shift in the relative energy actually *increases* the quenching. This is shown in Fig. 5a where we alter ε_S_1__ to bring either 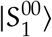 (ε_S_1__ = 14, 750cm^−1^) or 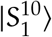 (ε_S_1__ = 13,600cm^−1^) into resonance with |TE^−^〉. In both cases 〈τ_ex_〉 drops by around 50% to roughly 300ps. Within the smaller range we find that changes in the energy of ε_S_1__ = 14, 050 ± 300cm^−1^, i.e. within the error bar of the reported value, the largest decrease is by about 25%. For ε_S_1__ > 18, 000cm^−i^ the quenching disappears completely (〈τ_ex_〉 → Γ_chl_ = 4ns), as has been previously reported [45]. This is simply because there is no energetic overlap between the two sub-systems and energy transfer between them is impossible by construction.

**Figure 5:**
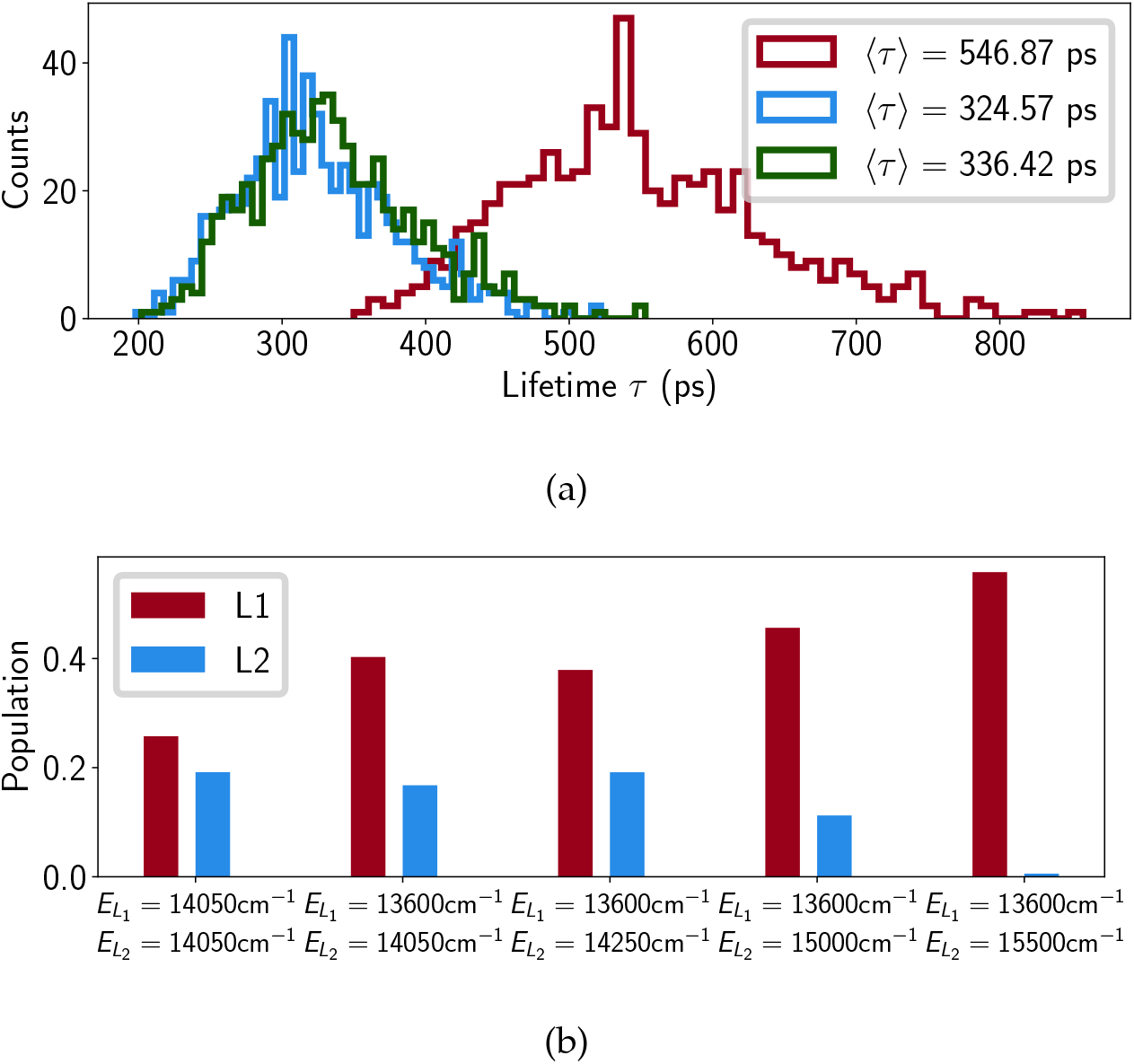
(a) Lifetime histograms with E_S_1__ set to 14050cm^−1^(red), 13600cm^−1^(blue) and 14750 cm^−1^(green), and (b) average populations on L1 and L2 at τ, where E_L_i__ denotes the 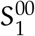 energy assigned to Lut i.

Although we initially excluded Lut2 we then put it back in the model, assuming the same transition charges and, initially, the same excitation energy as Lut1. The binding pocket of Lut2 (superficially) mirrors that of Lut1 with weak but significant couplings to Chls *a*603 and *a*604 which participate in several excitonic states between |TE^−^〉 and |TE^+^〉. This leads to a 40% decrease in lifetime relative to the Lut1-only model. Fig. 5b shows the excitation population on the Lut ground state at time t = τ_ex_ and we see that when 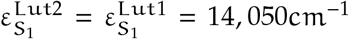 Lut2 is almost as effective a quencher as Lut1. This is contrary to the observed features of NPQ and the known properties of Lut2. The initial TA measurements that lead to the proposal of a Lut-mediated NPQ identified Lut1 as the sole quencher [23]. This is likely because Lut2 has a significantly distorted electronic structure relative to that in solution. The S_2_ excitation energy of Lut 2, 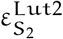, is significantly lower than 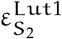 [58] and if this shift is caused by twisting of the backbone then it is likely accompanied by a concomitant upward-shift in 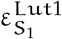 [59]. In fact, recent ultra-broadband 2D measurements on LHCII identified a dark state (termed S_x_), lying above the Chl a Q_y_ band, which belongs exclusively to Lut2 [60]. This is most likely a strongly-distorted S_1_. Fig. 5b shows that quenching by Lut2 can be completely abolished if we introduce some energetic asymmetry between Lut1 and Lut2. The shifts are not actually that large with 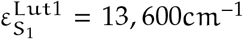 being almost within the error bars of the measured value of 14, 050 ± 300cm^−1^ and 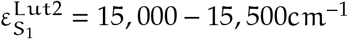 being roughly in the region of the proposed S_X_ state. It is important to note that this is not a rigorous analysis, which would require independent, *in situ* parameterization of Lut1 and Lut2. However, it points to an energetic sensitivity in the quenching pathway(s) that was absent in previous models.

## 3 Discussion

The essence of the quenching mechanism investigated here (and previously proposed [43]) is that trivial geometric modulation of Chl-Lut coupling is sufficient to drive the system behind quenched and unquenched states. This appears to be incorrect as the complex does not possess the conformational flexibility to induce significant changes in the coupling. We are not saying that the different minima do not represent functional states or that carotenoids are not involved in quenching, merely that our model does not capture its key features. There are several possible **quenching scenarios** that can be discussed.

### 3.1 NPQ may involve modulation of the properties of S_1_

The point of this study was to try and cast this problem in terms of experimental parameters, specifically the S_1_ energy, vibronic structure and relaxation dynamics. While the TA fits seem reliable, the data is for Lut in pyridine and obviously there is a question of whether this can be applied to Lut in protein. Actually, the default value of ε_S_1__ = 14, 050cm^−1^ was taken from NIR measurements on Lut in LHCII, although the error bars are quite big (±300cm^−1^) and the study did not compare quenched and unquenched configurations [46]. An earlier NPQ model proposed that quenching was induced by bringing the Chl Q_y_ band and S_1_ into resonance [25], which would require either 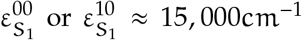. Balevičius Jr. and Duffy recently provided a very general physical argument as to why fine tuning of this energy gap cannot modulate quenching [52] and showed that significant quenching is possible for large energy gaps, even if the quenching state lies above the donor state. S_1_ has to be quite far above the Q_y_ band to abolish quenching, as was recently proposed by Lapillo et al. [45]. They reported a sharp dependence of the EET rate (and overall quenching) on the energy gap when ε_S_1__ ≈ 18, 000cm^−1^. This is simply because at this point the Q_y_ band coincides with the steep red edge of the S_1_ line-shape. However, as they point out, ε_S_1__ is not a free parameter and a protein-induced blue-shift of 14, 050 → 18, 000cm^−1^ (712 → 555nm) would be quite large. Saccon et al. recently performed TA measurements on quenched LHCII immobilised in polyacrylamide gel (a model for NPQ) [61] and found that linear excitation of Lut (i.e. via S_2_) produces the usual S_1_ → S_n_ ESA at ε_S_n__ - ε_S_1__ ≈ 18, 500cm^−1^ (540nm). This is reasonably close to the value in pyridine, ε_S_n__ - ε_S_1__ ≈ 17, 900cm^−1^ (558nm). Of course this is an indirect measurement and a massive shift in ε_S_1__ could be hidden by a correlated shift in ε_S_n__. However, this would have a significant affect on the ESA formation and decay kinetics which is not observed.

### 3.2 NPQ may involve non-Coulomb interactions and/or non-adiabatic inter-molecular states

For the planar geometry of Lut the published transition atomic charges [44] yield a dipole moment of |μ_S_1__| ≈ 2D which, although small, should certainly be detectable (|μ_Q_y__| ≈ 3 - 5D [62]). The fact that it is not implies that the amplitude of the S_1_ transition density is significantly over-estimated and therefore so are the Coulomb couplings, Q_y_ → S_1_ transfer rates, and overall level of quenching. In fact, given that there appears to be insufficient conformational flexibility in LHCII to switch this Coulomb-mediated quenching off, it may be purely an artefact. The reason that it was initially considered promising was that it qualitatively matched the NPQ scheme proposed by Ruban et al. in 2007, based on TA measurements of quenched LHCII aggregates [23]. The role of S_1_ was implied by global target analysis of the kinetics rather than any visually detectable S_1_ signal and so must be treated with caution. *Direct* observation of S_1_-mediated quenching was later reported for the cyanobacterial *High light inducible proteins* (Hlips) which are ancestors of LHCII [63]. Hlips are perpetually quenched by ≈ 2ps (hence observable) EET from a small pool of BChl a to β-carotene in one of the central binding pockets that are analogous to L1 and L2 in LHCII. More recently, sub-picosecond EET to Lut1 was directly observed in LHCII via ultra-fast 2D spectroscopy [58]. In both the EET is much faster than predicted by this or any of the previous models and it is difficult to see how such fast transfer could be Coulomb-mediated and be in any way switchable or involve an optically-forbidden transition. Cignoni et al. provide a possible answer via a detailed QM/MM study of CP29 in which short-range interactions (exchange, overlap, etc.) were found to make large contributions to the Chl-Cart couplings [64]. These are naturally far more sensitive to minor conformational changes than the long-range Coulomb interactions.

The picture gets even more complicated when one considers quenched LHCII in gel. It was recently shown that excitation of Q_y_ results in the immediate appearance of a large-amplitude positive peak at 19, 417cm^−1^ (515nm) which we’ll label A_515_ [61]. This is not merely a shifted S_1_ as direct excitation of Lut gave the usual S_1_ ESA at 18, 500cm^−1^ (540nm), although A_515_ is detectable at later times and may simply be initially hidden by S_1_. This suggests that S_2_ → S_1_ and S_2_ → A_515_ are competing pathways. A_515_ is in the region of the S* signal which some people suggest is either a distorted S_1_ or a dipole-forbidden singlet electronic state lying below S_1_ [65]. That argument aside, since the Chl ESA is typically flat and featureless, it seems reasonable to assume that A_515_ is associated with the Cars, although it is difficult to assign it solely to Lut1. A_515_ is independent of whether it is Chl *a* or Chl *b* that is excited and the GSB bands in the S_2_ region (< 500nm) looks very different to the classic S_2_ GSB. This all suggests some type of delocalized quenching pathway that involves several Carts and possibly even some non-adiabatic intermolecular states not accessible simply by exciting S_2_. This is exactly the picture emerging from the elaborate QM/MM models of CP29 being reported by Mennucci et al [45, 66].

### 3.3 Quenching requires hydrophobic mismatch and aggregation

It is possible that the conformational switch cannot be revealed by simulating a single LHCII monomer/trimer in a lipid bilayer. *In vitro* quenching is induced by low detergent concentration which in solution leads to aggregation. LHCII aggregates are the original model system for studying NPQ [20] and there is compelling evidence that some form of aggregation or clustering in the membrane is part of the *in vivo* mechanism [67]. Key to this is PsbS, with over-expression observed to enhance LHCII clustering and its absence frustrating it [68]. Recent simulations show that LHCII’s lumen-exposed side is covered in titratable residues with protonation causing an unfolding of a specific region implicated in protein-protein interactions [69]. Other studies have shown that it is capable of interacting with the minor PSII antenna complexes [70], possibly helping LHCII to partially detach from the reaction centre complex and form the quenching clusters. It has also been suggested that active PsbS has an affinity for certain lipids, altering local membrane composition and causing the hydrophobic mismatch that drives aggregation/clustering [71, 72]. It is therefore clear that any complete molecular model of NPQ will necessarily have to consider protein-protein interactions.

## 4 Methods

### 4.1 TA measurements of Lutein in Pyridine

All transient absorption data were measured with a spectrometer described in detail in ref [61]. Lutein (Sigma Aldrich) was dissolved in spectroscopic grade pyridine to yield an optical density of 0.2 mm^−1^ at the absorption maximum. The sample was placed in a 2 mm path-length quartz cuvette equipped with a micro-stirrer to avoid sample degradation during measurement. The mutual polarization of the excitation and probe beams was set to the magic angle (54.7°) and excitation intensity was kept below 1014 photons pulse^−1^cm^−2^.

### 4.2 The Chlorophyll exciton manifold

Modelling of energy relaxation within the chlorophylls is carried out according to previous work [34, 49] and is describe in detail in Section B the Supporting Information. Briefly, for a single uncorrelated MD snapshot (at time point t_k_) the relevant system of Chl excited (Q_y_) states is determined by the usual spin-boson Hamiltonian,

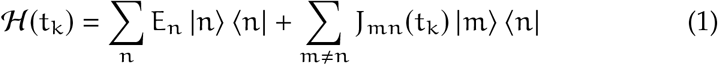

where {|m〉} is the ‘site’ basis of uncoupled single-molecule excitations, {E_m_} are the site (excitation) energies and {J_mn_(t_k_)} are the resonance couplings. {J_mn_(t_k_)} are calculated as the sum of pairwise Coulomb interactions between transition atomic charges, {q_∝_},

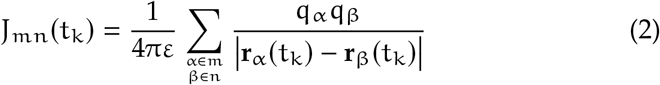

where ε = ε_r_ε_0_ = 2ε_0_. Both {E_n_} and q_∝_ are taken from Müh et al[73, 74]). Eq. 1 is then diagonalised to give the exciton basis,

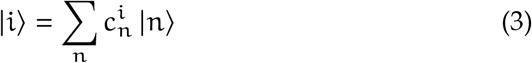

where 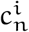 are the participation coefficients of each pigment state, |n〉, to a given exiciton state, |i〉. Site energies, oscillator strengths and couplings to Cart vibronic levels are also mixed. The exciton states are initially populated according to their oscillator strengths and relaxation is modelled using the modified Redfield approach outlined in ref. [34]. The population relaxation rates are given by,

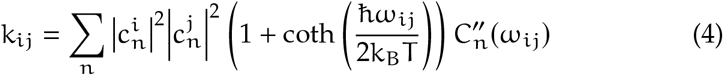

where 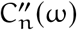 is the spectral density of bath-induced site energy fluctuations and ω_ij_ is the gap between the zero-phonon lines of excitons i and j. The ansatz spectral density from ref. [34] is assumed throughout. For a single uncorrelated snapshot along a trajectory (at time-point t_k_) the instantaneous LA and FL spectra are given by

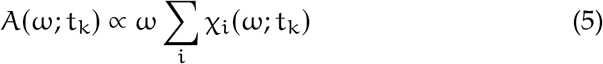

and,

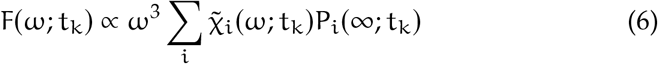

where {χ_i_(ω, t_k_)} and 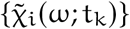 are the instantaneous LA and FL lineshapes respectively and {P_i_(∞; t_k_)} are the steady state populations of the exciton states. The true LA and FL for a particular minima are given by averaging over a trajectory, {J_mn_(t_k_)}, and then again over several instances of Gaussian disorder in the Chl site energies, {E_m_}.

### 4.3 The Carotenoid vibronic subsystem

The full VERA[53, 54] Hamiltonian is,

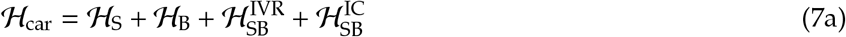

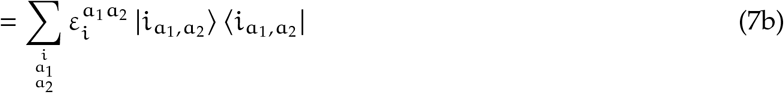

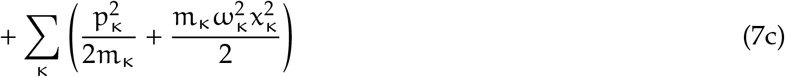

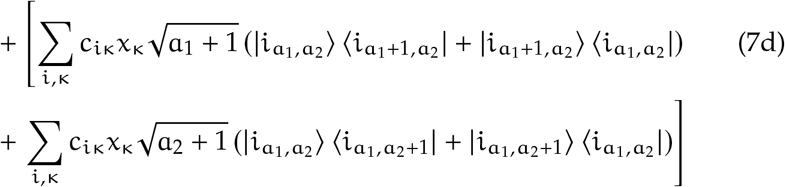

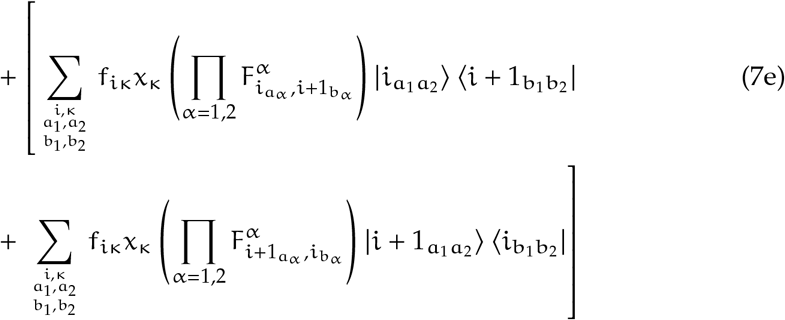

Eqn. (7b) is the Hamiltonian of the system 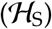 of uncoupled vibronic levels, {|i_a_1_ a_2__〉}, where

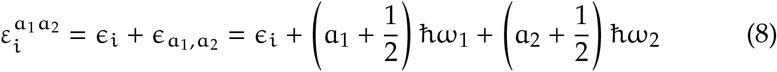

is the sum of the electronic, ∊_i_ and vibrational energies. 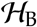 is the bath Hamiltonian which is composed of a large set of harmonic oscillators representing the non-optical modes of the the Cart itself plus librations, solvent modes, etc. We split this into two part, 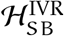 and 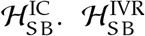 describes the bath-induced couplings between adjacent vibrational levels of the optical modes and are therefore responsible for vibrational relaxation on the electronic states. {c_iκ_} are the coupling constants and {x_κ_} the bath mode displacements. Energy (mostly) relaxes into the non-optical modes of the Cart and therefore reflect Intramolecular Vibrational Redistribution (IVR). Note, there is no population transfer between the two optically-coupled modes. 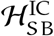 couples different electronic states and is therefore responsible for Interconversion (IC). It is characterized by coupling constants, f_iκ_, and the Franck-Condon (FC) overlaps,

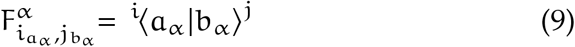

If we assume that the frequencies of the optical modes (ω_∝_) are independent of the electronic state (i.e. no Duschinsky rotations) then 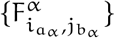 are entirely determine by their relative dimensionless displacements, 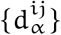. The relaxation dynamics are obtained by a second-order perturbative treatment of 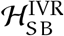 and 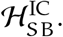 The resulting equations of motion are rather complicated and are listed in Section C of the Supporting Information. The various IVR, 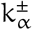, and IC, 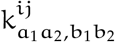, rate constants are defined in terms of Drude-type spectral density functions 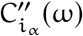 and C_i∝_,j_∝_(ω). We therefore have a large set of fitting parameters including electronic, {∊_i_} and vibrational, ∊_a_1_ a_2__, energies, modes frequencies, ω_∝_, mode displacements, 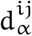, and the reorganization energies, λ_i_∝__, λ_i_∝_,j_∝__, and dephasing frequencies, γ_i_∝__, γ_i_∝_,j_∝__. Solving the dynamics yields a set of vibronic populations, 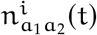, which are used to calculation the TA difference spectrum as a combination of ESA, GSB and stimulated emission (SE) components,

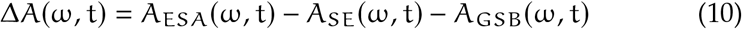

which are given by,

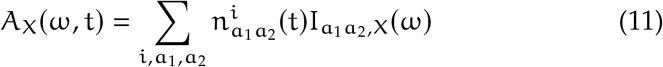

where I_a_1_ a_2__,x(ω) are FC-weighted Gaussian/Lorentzian lineshape functions that account for line-broadening.

### 4.4 Energy transfer between the Chlorophyll and Lutein subsystems

Having parameterized the subsystems separately we can now couple them via the calculated resonance couplings, 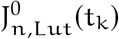. We make two assumptions. Firstly, since the inter-pigment couplings between the Chls and Lut is an order of magnitude smaller than the nearest-neighbour chlorophyll couplings (there is essentially no coupling between Lut1 and Lut2), we treat the Chl-Lut system as two weakly-interacting subsystems and assume that energy transfer proceeds incoherently [52]. Secondly, since there is almost no accumulation of vibronic population on the ground state (‘hot’ ground state), we do not explicitly include the Chl or Lut ground states in the dynamics. S_1_ can decay to higher vibrational levels on S_0_ but excitation proceeds from 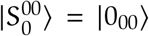. Essentially, we are assuming instantaneous IVR on the ground state. The couplings in the exciton basis are given by,

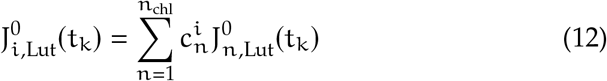

where 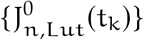 are the purely electronic Chl-Lut couplings. The rate of transfer from Chl exciton state |i〉 to Lut vibronic level 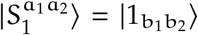 is given by the Fermi Golden Rule,

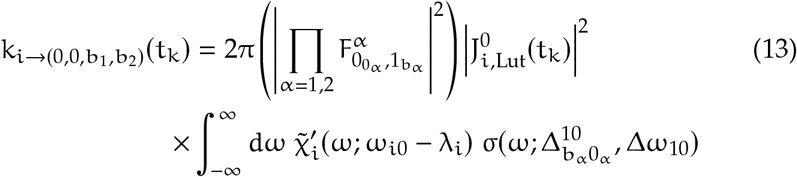

where 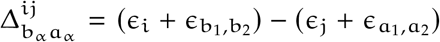 is the vibronic energy gap, 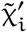 is the normalized excitonic fluorescence lineshape and 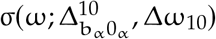 is the normalized vibronic Gaussian lineshape of width Δω_10_ = 1070cm^−1^ determined by the TA fit. The backward rate is similarly defined and Boltzmann factors are added to uphill rates to enforce the detailed balance condition.

## Acknowledgments

The authors thank Milan Durchan for assistance with transient absorption measurements. CG and CDPD thank the Biotechnology and Biological Sciences Research Council (BBSRC, BB/T000023/1). TW thanks the Chinese Scholarship Council for PhD funding. TP thanks the Czech Science Foundation (19-28323X) for financial support. VD would like to thank European Regional Development Fund and the Republic of Cyprus through the Research and Innovation Foundation (POST-DOC/0916/0049) for supporting the metadynamics studies of LHCII trimers.

## A VERA parameters for lutein

**Table 1:** The total parameter set for out vibronic model of Lut in pyridine, as described in the main text

|  |  |
| --- | --- |
| $\omega_{S_0}, \omega_{S_1}, \omega_{S_2}, \omega_{S_n}$ | $0 \text{ cm}^{-1}, 14,050 \text{ cm}^{-1}, 20300 \text{ cm}^{-1}, 31950 \text{ cm}^{-1}$ |
| $\omega_{C-C}, \omega_{C=C}$ | $1156 \text{ cm}^{-1}, 1523 \text{ cm}^{-1}$ |
| $\lambda_{S_0}, \lambda_{S_1}, \lambda_{S_2}$ | $15 \text{ cm}^{-1}, 100 \text{ cm}^{-1}, 150 \text{ cm}^{-1}$ |
| $\lambda_{S_0-S_1}$ | $31 \text{ cm}^{-1}$ |
| $\lambda_{S_1-S_2}$ | $860 \text{ cm}^{-1}$ |
| $\gamma_i, \gamma_{ij}$ | $163.6 \text{ fs}$ |
| $d_{C-C}^{S_0-S_1}, d_{C=C}^{S_0-S_1}$ | $0.82, 0.82$ |
| $d_{C-C}^{S_0-S_2}, d_{C=C}^{S_0-S_2}$ | $0.70, 0.84$ |
| $d_{C-C}^{S_1-S_2}, d_{C=C}^{S_1-S_2}$ | $0.80, 0.80$ |
| $d_{C-C}^{S_1-S_n}, d_{C=C}^{S_1-S_n}$ | $0.55, 0.0$ |
| $\Delta\omega_{S_0-S_1}, \Delta\omega_{S_0-S_2}, \Delta\omega_{S_1-S_n}$ | $1070 \text{ cm}^{-1}, 1190 \text{ cm}^{-1}, 1090 \text{ cm}^{-1}$ |
| $\frac{\vert \mu_{S_1-S_n}\vert ^2}{\vert \mu_{S_0-S_2}\vert ^2}$ | $1.22$ |
| $S_2$ Stokes shift | $150 \text{ cm}^{-1}$ |

## B Secular Redfield model of the Chl manifold

As outlined briefly in the main text we use secular Redfield theory for the modelling of energy relaxation on Chl exciton manifold. The starting point is the spectral density of energy gap fluctuations for the uncoupled Chls for which we assume the ansatz spectral density proposed by Novoderezhkin et al [34].

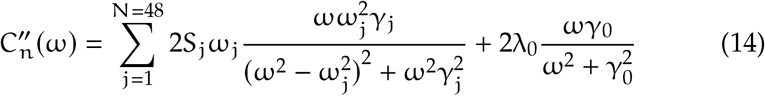

Here ω_i_ are the frequencies of the 48 under-damped modes, S_i_ are the associated Huang-Rhys factors and γ_i_ are the damping times. The overdamped (Drude) term is characterized by is own damping time, γ_0_, and a reorganization energy λ_0_. The effective reorganisation energy is given by

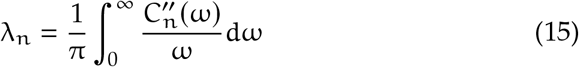

and the energy-gap correlation function (in the frequency domain) is given by the fluctuation-dissipation theorem,

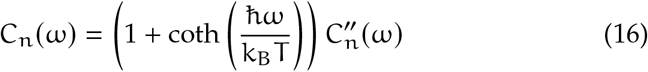

Switching to the exciton basis, the transition dipole moments, reorganization energies, relaxation rates and correlation functions (in the time domain) mix according to,

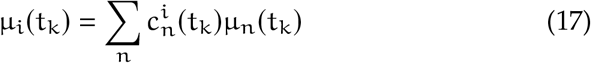

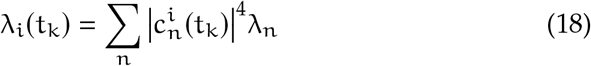

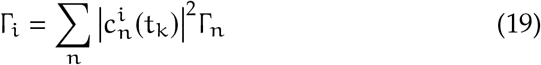

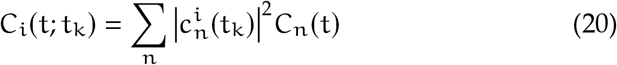

with the participation coefficients 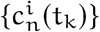 and uncoupled (site) transition dipoles, μ_n_(t_k_) varying from snapshot to snapshot. 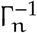 are the excitation lifetimes of the uncoupled Chls which are all assinged the typical experimental value of 4ns. The exciton line-broadening functions are expressed in terms of 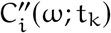

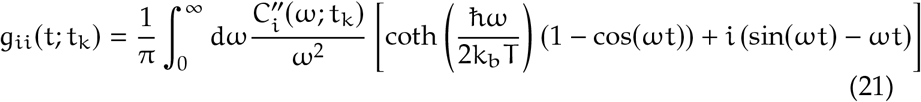

which in turn gives the instantaneous (snapshot) exciton LA,

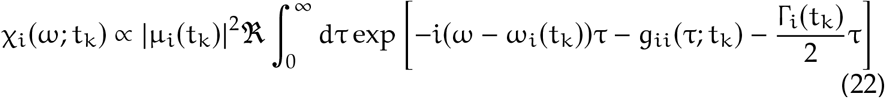

and FL,

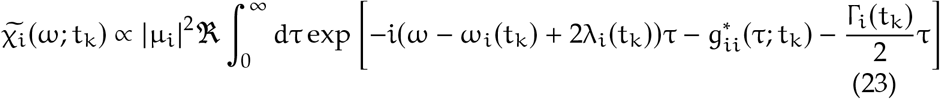

line-shapes respectively. These are used to calculate the LA and FL spectra as in the main text. Finally, the population dynamics, {P_i_(t; t_k_)}, of a single MD snap-shot are given by a set of Master Equations,

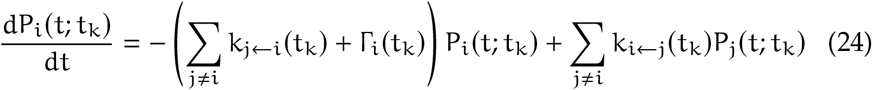

with the rate constants defined in the main text. Note we completely ignore the coherences as they are not relevant to the timescales being probed.

## C The Vibrational Energy Relaxation Approach (VERA) to the Lut dynamics interaction

We use the VERA approach developed by Balevičius et al. [53] the basic assumptions of which are discussed in the main text. The time-evolution of the vibronic populations are given by,

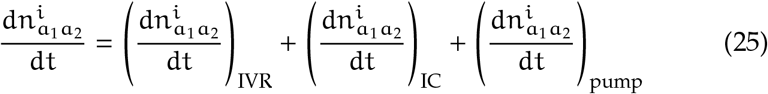

where the three terms correspond to the vibrational relaxation on the electronic states (IVR), interconversion (IC) between electronic states, and the initial resonant excitation by the pump pulse. The IVR is determined by

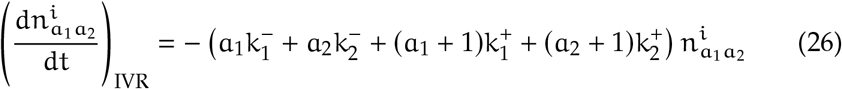

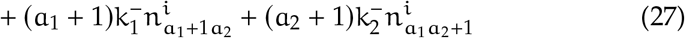

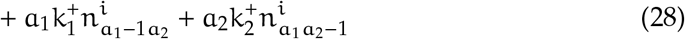

where the first line denotes loss of population to upper and lower vibrational states on mode 1 and 2 and the second set of four terms denote incoming population from those states. We define upward, k^+^, and downward, k^−^, vibrational relaxation rates as

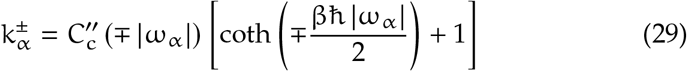

Note that in the harmonic approximation we neglect overtone (Δa_∝_ > a_∝_±1) transitions or couplings between the two modes. For the IC dynamics we have

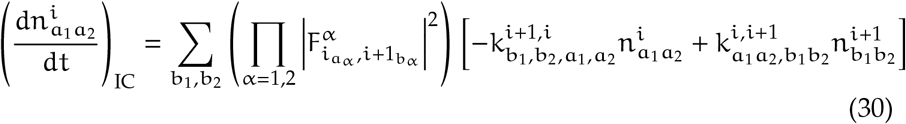

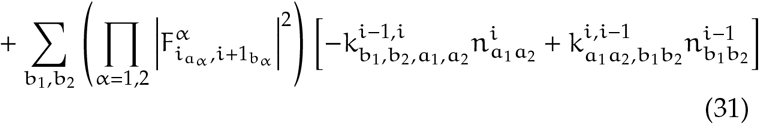

where the first pair of terms describe the upward and downward transitions between |i_a_1_a_2__) ↔ |i + 1_b_1_b_2__) respectively and the second pair describe the upward and downward transitions between |i_a_1_a_2__) ↔ |i - 1_b_1_b_2__) respectively. Here the rate constants 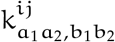, are defined as

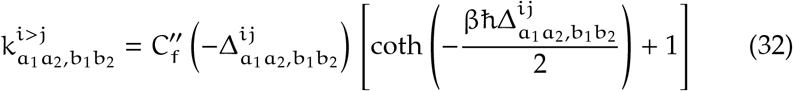

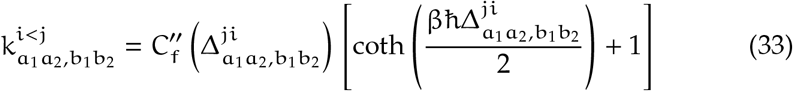

with 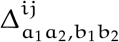 defined as in the main text.

